# Extensions of mean-field approximations for environmentally-transmitted pathogen networks

**DOI:** 10.1101/2022.09.13.507803

**Authors:** Kale Davies, Suzanne Lenhart, Judy Day, Alun L. Lloyd, Cristina Lanzas

## Abstract

Many pathogens spread via environmental transmission, without requiring host-to-host direct contact. While models for environmental transmission exist, many are simply constructed intuitively with structures analogous to standard models for direct transmission. As model insights are generally sensitive to the underlying model assumptions, it is important that we are able understand the details and consequences of these assumptions. We construct a simple network model for an environmentally-transmitted pathogen and rigorously derive systems of ordinary differential equations (ODEs) based on different assumptions. We explore two key assumptions, namely homogeneity and independence, and demonstrate that relaxing these assumptions can lead to more accurate ODE approximations. We compare these ODE models to a stochastic implementation of the network model over a variety of parameters and network structures, demonstrating that with fewer restrictive assumptions we are able to achieve higher accuracy in our approximations and highlighting more precisely the errors produced by each assumption. We show that less restrictive assumptions lead to more complicated systems of ODEs and the potential for unstable solutions. Due to the rigour of our derivation, we are able to identify the reason behind these errors and propose potential resolutions.

## 1 Introduction

Infectious diseases spread through a population via different modes of transmission, including direct contact, sexual contact, vectorborne, airborne, and environment-to-host transmission. Many infectious diseases that cause high morbidity and mortality are able to infect via environments such as surfaces, soil or water without direct contact between hosts [5, 13, 14, 35, 36]. To explicitly address environmental exposure pathways, models for environmentally-transmitted pathogens have taken an analogous approach to their direct transmission predecessors, namely through the development and analysis of deterministic ordinary differential equation (ODE) models [6, 28, 37, 47] or stochastic individual-based models [7, 45, 54]. In both cases, a key element of environmental transmission models is that the infectious state and the pathogen population are explicitly modelled.

The majority of ODE models for environmental transmission are mean-field models in which both the infected population and the pathogen population are assumed to be independent and randomly-mixed. Given that environmental transmission problems depend on host population structure, pathogen dynamics, and spatial scale of the modeled system, these assumptions are often not appropriate [16, 32, 52]. Within the direct transmission literature, progress has been made in resolving the issues with mean-field models [43], including pair approximations via moment closure [23, 51], clustering of nodes with similar properties [29, 15] and quantifying approximation errors [12, 4]. These same improvements has not yet reached the environmental transmission literature, except for in some rare cases [55]. In this paper, we apply some of the existing approaches for direct transmission to develop ODE models with less restrictive assumptions in the context of environmentally transmitted infections.

In contrast to the mean-field models, individual-based models [44, 53] consider all individuals within a system, explicitly modelling their local interactions, usually with stochastic features. While this allows the independence and homogeneity assumptions to be completely relaxed, it does come with some setbacks. A complex individual-based computational simulation based on rules and individual attributes can be quite resistant to mathematical analysis or sometimes even to experimental numerical intuition [1, 31]. Further, large or detailed populations [46, 42] typically result in individual-based models that are computationally expensive and sensitive to parameter changes and initial conditions, further increasing the difficulty in analysis.

Individual-based network models have been used in the analysis of directly transmitted infections for a number of purposes [2, 10, 19, 20, 25, 34, 41]. Using a network, we can describe a population of individuals, uniquely describing individual characteristics and interactions between individuals—avoiding the assumption of random mixing. Further, by constructing the network to be a continuous-time Markov chain, it can be implemented as a stochastic simulation while also being used as a rigorous basis to derive compartmental ODE models [22, 26, 49, 51]. Environmentally-transmitted pathogen dynamics inherently depend on the environmental processes and spatial scale of the problem, all of which can be taken into account in the network structure. These benefits make networks a suitable approach to characterize transmission for environmentally-transmitted pathogens.

The primary goal of this work is to present a clear and rigorous derivation of systems of ODEs for approximating the average infected population of a stochastic network model undergoing an environmentally transmitted epidemic. Such a derivation has not been demonstrated for environmentally transmitted infections and presents some unique situations not seen in the analogous direct transmission network. The specific epidemic dynamics of our network are inspired by hospital-acquired infections, in which the nodes represent patients in hospital rooms and pathogen can be transferred between rooms by healthcare workers [3, 9]. Despite this, the techniques used throughout this paper can be applied broadly to other environmental transmission dynamics. In total, we derive four different approximation models, using assumptions of statistical independence, homogeneous mixing and moment closure [11, 22, 27, 30, 49, 51, 48, 33]. By assuming independence and homogeneous mixing, our most restrictive set of assumptions, we derive a mean-field model, a form frequently used in the environmental transmission modelling literature [6, 8, 18]. We show that in deriving this model, almost all network structure is lost. Each of our other ODE models (Models B, C, D) aim to improve upon Model A by enforcing less restrictive assumptions at the expense of requiring a larger system of ODEs. By making fewer assumptions we attempt to minimize the error between the ODE models and the true behaviour of the stochastic network model.

The secondary goal of the paper is to give a qualitative analysis of the accuracy of the approximation models based on the assumptions implemented in each derivation. This is done by solving the systems numerically and comparing the approximations to simulated realizations of our network model. We show that by removing both the independence and homogeneity assumptions (Model D) the average network behaviour can be approximated with much greater accuracy for some parameter choices, but that unstable behaviour of the ODE model prevents this for other parameter choices. Finally, we note our goal of determining the exact impact of each assumption is difficult as the validity of each assumption is inherently tied to the chosen environmental transmission dynamics. We support our claims primarily by considering a variety of parameter sets and network structures, and by observing errors and improvements that are consistent with the expected consequences of the assumptions.

## 2 Defining the stochastic network model

### 2.1 Network description

Consider a network of *N* nodes with a set of edges defined by adjacency matrix *G* with entries *g*_*ij*_ = 0 if there is no edge and *g*_*ij*_ = 1 if there is an edge between nodes *i* and *j*. Our network is undirected, hence *g*_*ij*_ = *g*_*ji*_ for all nodes *i* and *j*, although results should hold regardless of this condition. In the case where *g*_*ij*_ = 1 we refer to nodes *i* and *j* as being adjacent. We will be considering nodes that represent both a position in space and unique individuals. In practice, this is a suitable way to represent patients in a hospital setting, with nodes representing beds or rooms, each of which will only be occupied by exactly one patient at all times. As such, nodes have two components to their states: infection state and pathogen state. The infection state of a node can be susceptible or infected, representing the status of the individual associated with that node. The pathogen state of a node represents the amount of pathogen in that location in space. The units of the pathogen state are flexible (could be colony-forming units, number of parasites, proportion of space pathogen occupies or a general hazard level [47, 50]) and will generally be based on the specific host-pathogen system, application and access to data.

The system has 5 different events that allow the network nodes to change states: 1) susceptible individuals can become **infected** by exposure to the pathogen on the same node; 2) infected individuals can **recover** to the susceptible state; 3) infected individuals can **shed** the pathogen, increasing the pathogen state on the same node; 4) pathogen can **decay**, decreasing the pathogen state; and 5) pathogen can **move** from one node to an adjacent node. These events, the rates at which they occur and their exact changes to the network are given more precisely in Table 1. Note that some events require interactions, either between nodes (movement) or between the pathogen and infection state of a single node (infection, shedding), while other events can occur without an interaction (recovery, decay).

**Table 1:**
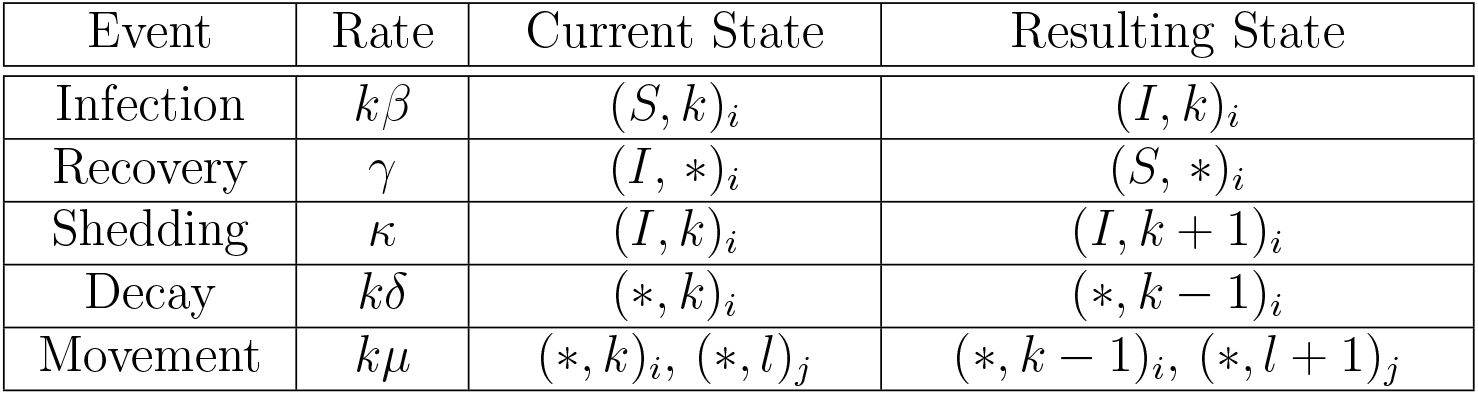
A list of the different events, the rate at which they occur, the required state of the system and the resulting state after the event takes place. The notation (*X, k*)_*i*_ denotes node *i* having infection state *X* and pathogen state *k*. If an event neither requires nor changes either the infection or pathogen state of a node, then we denote this with the ∗ symbol. Movement events can only occur if nodes *i* and *j* are adjacent.

Events cause changes in the state of the system which can be measured on a number of levels. We have already defined both infection state and pathogen state. It is also convenient to define the node state that encompasses both of these two components. A given node will have its node state change if the infection or pathogen state is changed by an event. We then define the network state as the union of all node states. As such, the network state will change if any event occurs. We can use the events in Table 1 to describe the transition rates from a given network state to any other possible state. As these transitions depend only on the current network state, the network dynamics can be represented by a continuous-time Markov chain (CTMC).

We define the random variables *I*^(*i*)^ and *P*^(*i*)^ such that they represent the infected state of node *i* and the pathogen state at node *i* respectively. As such, we can write the probability of node *i* having infection state *X* and pathogen state *k* as

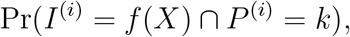

where the element *X* ∈ {*S, I*} represents the infection state, *k* ∈ {0, 1, 2, …} is the pathogen state and *f* : {*S, I*} → { 0, 1} is a function that transforms the state *X* into a numerical value such that

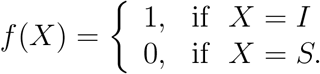

Note that we also use the random variable *S*^(*i*)^ where convenient. We define this variable such that *S*^(*i*)^ + *I*^(*i*)^ = 1.

### 2.2 Measuring average behaviour

Ultimately our objective is to measure the average behaviour of the network model, specifically the expected number of infected individuals and the average pathogen state of the network at all times. Ideally we would use the exact system, but in almost all cases it is impractical to deal with directly due to its size and complexity. Here, we outline the difficulties in obtaining an exact value for the expected number of infected individuals and outline two possible approaches for approximating and estimating this quantity. Namely, we will look at the exact system, the simulated system and the approximate system. The relationships between these systems and the network are presented in Figure 1.

**Figure 1:**
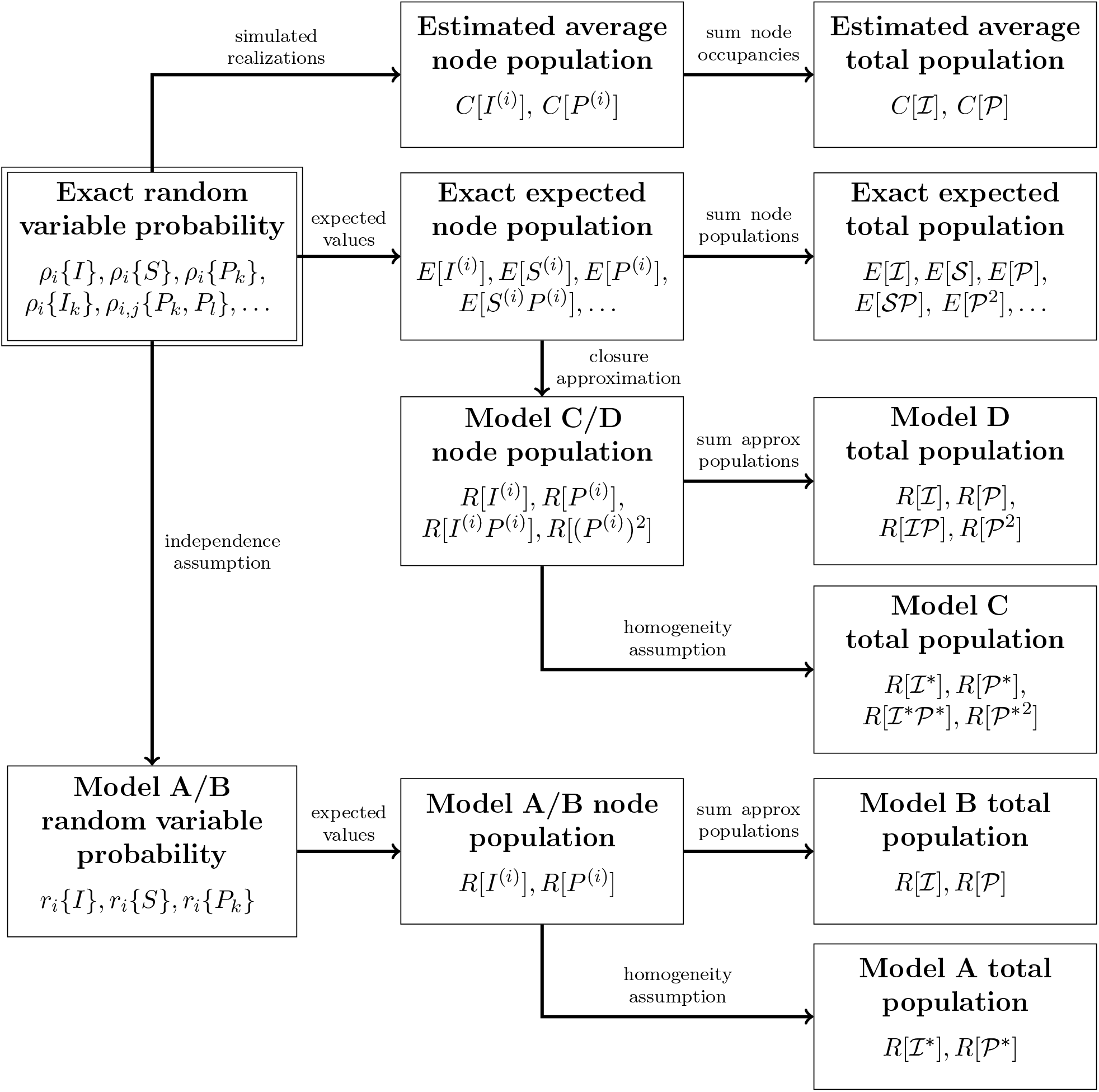
A visualization of the relationship of different systems corresponding to the network. Each box represents a different way of describing the network’s population, including terminology and notation used in each case. Every approximation starts from our most general and informative system, describing the exact probability of each random variable in the system (top left). Following any arrow will reduce the size of the system, making it easier to solve, however this comes at the expense of information. In some cases, this means that we are unable to describe specific node behaviour but are still able to accurately describe the population behaviour, while in other cases we are making approximations and fundamentally changing the structure and dynamics of the original system. All notation presented here is precisely defined in Section 2.2.

#### 2.2.1 Exact system

We begin by considering the exact system, given by each box in Figure 1 that begins with “Exact”. Theoretically, the expected node populations can be obtained via the Kolmogorov equations (known in some fields as the Master equations) from the CTMC.

First, we define the following notation for the probability of a node being infected and in pathogen state *k*

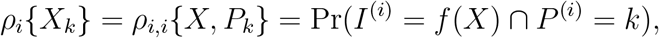

where *X* ∈ {*S, I*} represents the infection state and *k* ∈ 0, 1, 2, … is the pathogen state. Using this notation, we may also write the probability of a node being infected regardless of its pathogen state, *ρ*_*i*_{*I*} or the probability of a node being in pathogen state *k* regardless of its infection state, *ρ*_*i*_{*P*_*k*_}. We can also denote more complex states, for example node *I* being susceptible and in pathogen state *k* while node *j* is in pathogen state *l* regardless of its infection state, *ρ*_*i,j*_{*S*_*k*_, *P*_*l*_} = *ρ*_*i,i,j*_{*S, P*_*k*_, *P*_*l*_}.

In order to set up our CTMC, we define ***ρ***(*t*) the network state vector. Each entry of ***ρ*** gives the probability of being in its corresponding network state at time *t*, that is, each entry is the probability of a given configuration of the entire network. As the network dynamics can be represented by a CTMC, there exists a *Q* matrix such that the exact probability distribution of each network state is given by

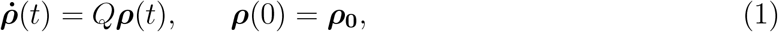

for some initial condition ***ρ***_**0**_ [40]. The system of differential equations given by equation (1) are the Kolmogorov forward equations. The entries of *Q* are the transition rates between pairs of network state. As transition rates are independent of the current time, the entries of *Q* will be constant. This equation gives us a fundamental starting point for describing the average behaviour of the system, however writing the explicit components of *Q* is unnecessary. We will briefly address this again as we begin to derive the system in Section 3.

If we can solve the system of differential equations (1) simultaneously, we obtain the exact probabilities of being in each network state. From this we can extract the node states and the infection and pathogen states of each node. The number of equations in this system is given by

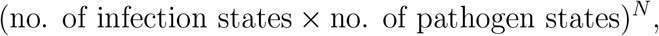

where *N* is the number of nodes in the system. This system is unwieldy even with very few pathogen states and very few nodes. As such, it is not practical to solve the system using the Kolomogorov forward equations (equation (1)) directly.

It should also be noted that the number of pathogen states is technically unbounded, meaning that *ρ* is an infinite vector and *Q* is an infinite matrix. While this could cause some issues for our derivation, we note that so long as the pathogen decay rate *δ* is non-zero, the expected value of the pathogen will have an upper bound. As we will eventually rewrite our systems in terms of expected values, this boundedness will ensure that there are no issues with our derivation.

We can consider a system of reduced size by instead considering expectations. The following equations give the expected number of infected individuals at node *i* and the expected pathogen state at node *i*

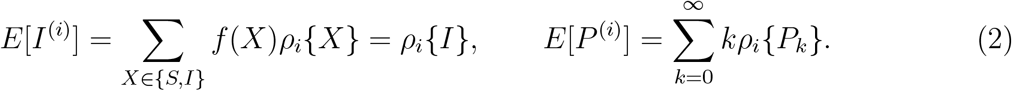

If we derive differential equations for these variables and solve them, they give us the exact average node occupancies. Similarly, if we sum these expressions over all nodes, we obtain the expected average populations over the network

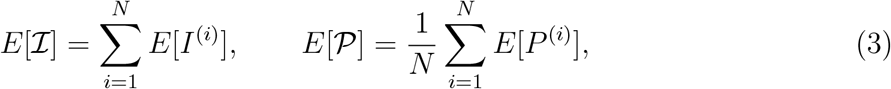

where ℐ is a random variable for the number of infected individuals in the network, and 𝒫 is a random variable for the average pathogen state per node across the network.

Unfortunately, in most cases we either cannot derive exact rate of change equations or we are unable to form a closed system without increasing the number of variables to the point where it is impractical to solve. Instead, there are two common approaches to obtain the expected populations: (1) simulating the system and averaging over many realizations, or (2) developing approximating equations based on the exact equations.

#### 2.2.2 Simulated system

The simulated system is given in each box in Figure 1 that begins with “Estimated”. As the network model is a CTMC, we can simulate the process via the Gillespie algorithm [17, 24] using the following steps:

1. calculate the total rate *λ*, by summing the rates of all possible events that can occur from the current state,
2. generate a time step that is exponentially distributed with parameter 1*/λ*,
3. randomly select an event to implement, proportional to the rate at which each event can occur,
4. implement the event and the time step and repeat the process.

Each event that can occur contributes to the total rate *λ*. Specifically, *λ* is the sum of the nondiagonal elements of the column of *Q* corresponding to the current state. The requirements for each event can be observed in Table 1.

At each point in time, we average over the set of simulated realizations in order to obtain the estimated average node occupancies *C*[*I*^(*i*)^] and *C*[*P*^(*i*)^] at that time. Similarly, we sum over each node to obtain the estimated average population *C*[ℐ] and *C*[𝒫]. The larger the number of realizations, the more accurate each of these estimates become.

Unfortunately, simulating many realizations is computationally expensive. Depending on the choice of parameters, the number of required realizations and the structure of the network, it can take hours or days to obtain the estimates. A practical alternative to resolve these issues is using approximate systems.

#### 2.2.3 Approximate systems

The primary issue with the exact equations is that they cannot be reduced except for a few special cases. To make these equations manageable, it is common to make approximations that reduce the size of the system, albeit at the expense of accuracy. The different approximate systems, each under different assumptions, are given by each box in Figure 1 that begins with “Model”. Our goal here is to highlight the assumptions of each model, which may give us necessary insight when interpreting the results of our models.

We define a new set of variables *r*_*i*_{∗} to denote the approximate probabilities, with each variable being the approximation of its exact system counterpart *ρ*_*i*_{∗}. The new set of variables is defined to have both the same initial condition and structure as the exact system, but with a assumption that reduces the total number of variables in the system [11, 26, 49]. For example, the approximate system might use

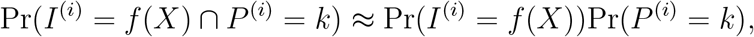

so where a variable such as *ρ*_*i*_{*I*_*k*_} exists in the exact system, it would be replaced by *r*_*i*_{*I*} *r*_*i*_{*P*_*k*_} in the approximate system. The implementation of this approximation yields a system of differential equations in which the node states are no longer considered explicitly, with all terms being reduced to the probability of an infection or a pathogen state, resulting in a much smaller system of differential equations. However, such an independence assumption implies that the covariance of the random variables *I*^(*i*)^ and *P*^(*i*)^ is zero. Despite the unrealistic nature of this assumption, it is frequently implicitly used, particularly in mathematical epidemiology in the context of Susceptible-Infected-Recovered (SIR) compartmental models.

Even upon making this assumption, the system of differential equations is still unbounded as the pathogen state is unbounded, meaning the system cannot be solved directly. To reduce the system further, we rewrite it in terms of approximate expected values, *R*[∗]. We define the following approximate expected values such that they are analogous to the expected values of the exact system (equation (2))

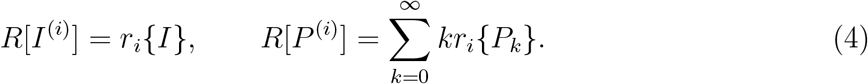

We will also see cases where applying independence assumptions to the probabilities does not allow us to derive equation for the approximate expected values. In these cases, we can write a system of differential equation for the expected value of the exact system and implement a different approximation known as moment closure [22, 27, 51]. In doing so, assumptions are made about the distribution of the random variables, allowing us to rewrite the system in terms of lower order moments. We will discuss this in more detail in Section 4.

If we can solve the system of differential equations for *R*[*I*^(*i*)^] and *R*[*P*^(*i*)^], then we can calculate the approximate expected population over the whole network. Specifically, this is done by writing approximate expected values for our population random variables which are analogous to the expected values of the exact system (equation (3))

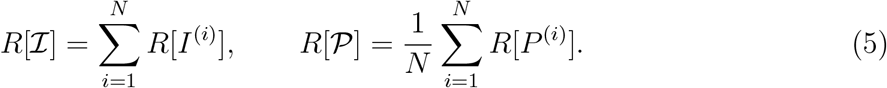

In cases where the system of differential equations for *R*[*I*^(*i*)^] and *R*[*P*^(*i*)^] can be obtained, we still require a differential equation for each node. Not only is the system difficult to analyse directly, but it is computationally expensive if the number of nodes is large. Ideally, we would obtain a closed system of differential equations for *R*[ℐ] and *R*[𝒫], however in general this is impossible without making one more assumption.

To further simplify the system of differential equations, we make an assumption of homogeneity among nodes. This assumption is also commonly referred to as ‘homogeneous mixing’ or a ‘well-mixed’ assumption. Assuming homogeneity spreads the population equally among all nodes, meaning that if only one node was infected in the network, the system would instead consider all nodes as being 1/*N*th infected. To represent this change, we stop considering the population random variables ℐ and 𝒫, and instead consider the two analogous homogeneous population random variables ℐ^*∗*^ and 𝒫^∗^, for which *I*^(*i*)^ = *I*^(*j*)^ and *P*^(*i*)^ = *P*^(*j*)^ respectively, for all nodes *i, j*. These new random variables will leave all nodes indistinguishable from one another using the definitions

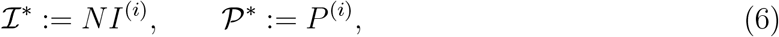

for all nodes *i*. This results in the following equations for the approximate expected values

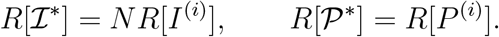

The homogeneity assumption removes most aspects of the network structure from the approximation. In networks with differing numbers of edges connected to each node, the movement rate would usually differ between nodes. As the model is no longer able to distinguish between nodes, we replace the movement rate *µ* with the average movement rate among all nodes. We call this the adjusted movement rate *µ*^∗^, which is given by the equation

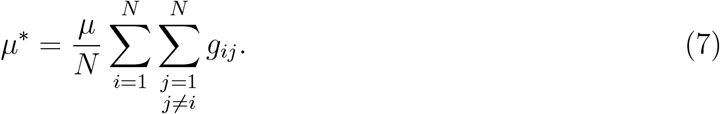

Noting that ∑ ∑*g*_*ij*_ is simply the number of non-zero entries of *G*, we can see that the network’s structure has been reduced to the number of edges in the network.

Just as with the simulated system, this approximation approach has a number of consequences. The primary trade-off here is the error introduced by the approximation versus the computational expense. The simplest system we will show in our work, Model A, is the most commonly used model in the literature and has little computational expense, but is often an extremely inaccurate approximation to the exact system. As approximations use less restrictive assumptions, the computational expense increases and more complex systems arise that are more difficult to solve analytically. Further, the assumptions used can have unpredictable consequences. Qualitatively we know that assumptions of independence generally cause models to overestimate the number of infected individuals [49], however it is difficult, if not impossible, to predict the error produced without simulating the system. We discuss these ideas with more concrete examples after we derive our models below.

For the remainder of this paper, we will derived approximate systems that aim to adequately approximate the expected occupancy for populations at a node or over the network. We will consider two approximations that use an independence assumption (Models A,B), while the other two systems bypass this and require a moment closure approximation (Models C,D). Similarly, only one of the two systems in each case apply a homogeneity assumption (Models A, C), while the other two systems calculate the approximate populations for each node and use summations to obtain the approximate population over the network (Models B, D). This will result in four different models overall. Once derived these models will be compared to each other and to the estimated average population calculated via the simulated system. This will allow us to better discuss the nature of the assumptions and their consequences.

## 3 Independence system

We derive differential equations that calculate the number of infected nodes in the network at any given time. We have a secondary objective of tracking the value of the pathogen. As our primary objective is the number of infected nodes, we first consider the exact change in the probability of an arbitrary node *i* being in an infected state

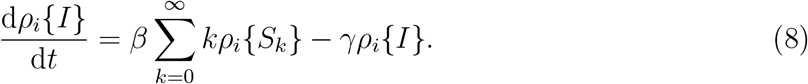

These equations can be obtained rigorously from equation (1) by summing over all states that included the *i*th node being infected [40], however it is generally not feasible to do this directly. Instead we intuitively construct these equations by considering the events that change the infectious state of a node: a node can leave an infected state via a recovery event, or a node can enter an infected state via an infection event. In the case of recovery, the event occurs at rate *γ* and requires a node to be in an infected state (resulting in the second term). In the case of infection, the event occurs at rate *β* per pathogen state, that is *βk*, and requires a node to be in a susceptible state and in pathogen state *k* (resulting in the first term).

Equation (8) cannot be solved without knowledge of *ρ*_*i*_{*S*_*k*_} and how it changes over time. We refer to *ρ*_*i*_{*S*_*k*_} as a ‘second order’ term, as it refers to two properties: the infection state of node *i* and the pathogen state of node *i*. In contrast, we refer to *ρ*_*i*_{*I*} as a ‘first order’ term, as it refers to only a single property, namely the infection state of node *i*. We make a simple approximation by considering only first order terms. To bypass this second order term, we apply the following approximation

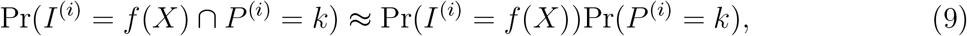

splitting it into the product of two first order terms. This expression assumes independence between the infection state and pathogen state of a given node. Such an assumption is only valid if there is no covariance between the two random variables.

To formalise this approximation, we describe a new system that represents our approximation to the true solution, under the specified set of assumptions. We define ***r*** such that it corresponds to ***ρ***; the *i*th entry of ***r*** is an approximation for the *i*th entry in ***ρ*** (*r*_*j*_{∗} is an approximation for *ρ*_*j*_{∗}, for all nodes *j*). We then define an approximate system with the following properties:

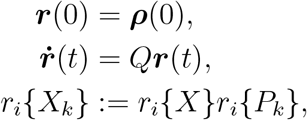

for all nodes *i, j* and all pathogen states *k*. These properties describe the approximate system as being identical to the exact system in terms of initial condition and structure, while defining some variables differently to express the desired approximation. This approximation corresponds to equation (9).

Using this new approximate system, we can write the approximate equation that is analogous to equation (8):

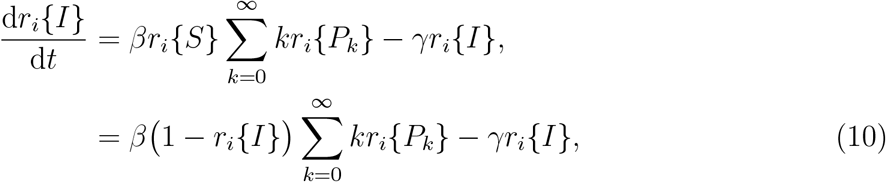

noting that *ρ*_*i*_{*S*} + *ρ*_*i*_{*I*} = 1 is a property of the exact system, and the corresponding property *r*_*i*_{*S*} + *r*_*i*_{*I*} = 1 continues to hold for the approximate system.

In order to solve equation (10), we also require a differential equation for the term *r*_*i*_{*P*_*k*_}. The derivation, explained below, results in the following exact equation:

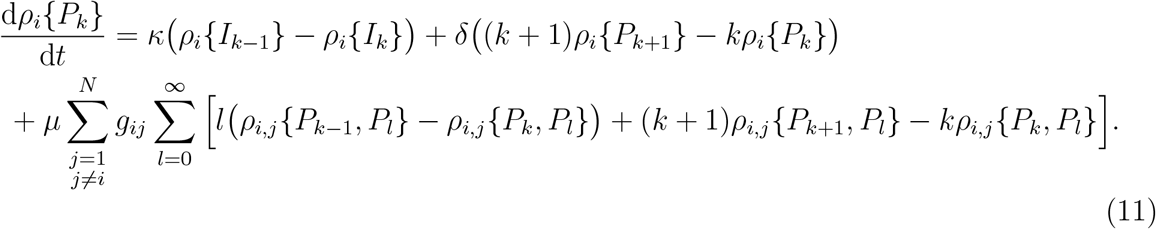

noting that at the pathogen state boundary *k* = 0, we will obtain slightly different equations (specifically, terms that would be *k* = −1 are removed).

Equation (11) can be derived using the same approach as equation (8), by intuitively considering possible changes to the pathogen state based on each event in the network. The shedding event occurs at rate *κ* and can either enter pathogen state *k* from state *k* − 1, or leave pathogen state *k* to state *k* + 1, so long as the node is in an infected state. The decay event occurs at rate *δ* per pathogen state and can either enter pathogen state *k* from state *k* + 1 (at rate *δ*(*k* + 1)) or leave pathogen state *k* to state *k* − 1 (at rate *δk*). Similarly, the movement event can cause a node to enter or leave pathogen state *k*, however there are four ways for this to happen. The four movement terms in order are: (1) node *i* entering pathogen state *k* after receiving from an adjacent node, (2) node *i* leaving pathogen state *k* after receiving from an adjacent node, (3) node *i* entering pathogen state *k* after giving to an adjacent node, and (4) node *i* leaving pathogen state *k* after giving to an adjacent node. The summations correspond to looking over all pathogen states and over all nodes (but only including adjacent nodes, that is *g*_*ij*_ = 1). Following similar logic we can write the rate of change equation for the probability of any given state (or sub-state) of the system. For the remainder of our derivations, we will be stating exact equations such as equations (8) and (11) without further discussion.

In order to rewrite equation (11) using only first order terms, we need to reduce the second order terms of the form *ρ*_*i,j*_{*P*_*k*_, *P*_*l*_}. While in general we can do this using an approximation, our specific system dynamics cause these terms to naturally break down later when calculating expected values. As such, it is sufficient for our purposes to rewrite equation (11) using the approximate system defined above

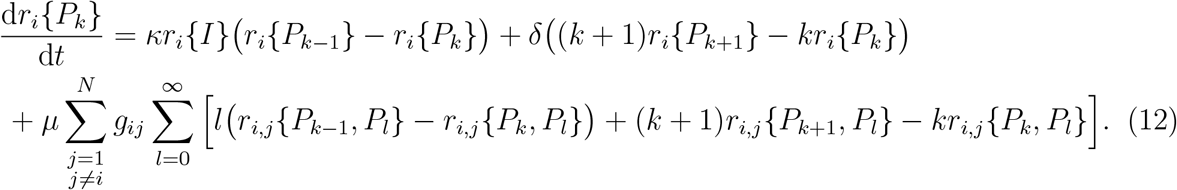

We now rewrite these approximate probability equations in terms of approximate expected values. We usLe equation (4) along with the total probability equations (namely *ρ*_*i*_{*S*} + *ρ*_*i*_{*I*} = 1 and ∑_*k*_*ρ*_*i*_{*P*_*k*_} = 1) to rewrite equation (10) as

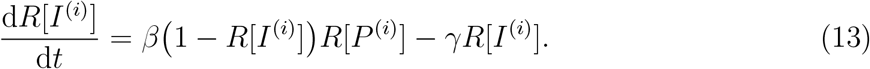

For equation (12), we must first multiply the equation by *k* (corresponding to the pathogen state) and then sum over all pathogen states. Doing this yields

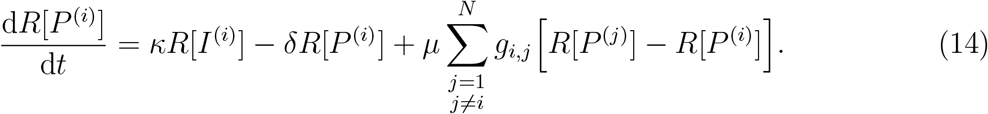

This approximate system has initial conditions

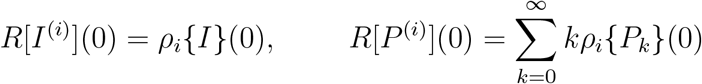

Equations (13) and (14) (with the given initial conditions) can be solved to give the approximate expected node populations.

Recall that our goal is to obtain expect values for the number of infected nodes ℐ and the average pathogen state over all nodes 𝒫. Upon solving equations (13) and (14) we can approximate the expectation of these random variables via equation (5).

Unfortunately, due to the non-linear term in equation (13), namely *R*[*I*^(*i*)^]*R*[*P*^(*i*)^], we can make no simplifications to this expression without making further assumptions. As such, we implement a homogeneity assumption, which will remove almost all spatial components from the model, considering both infected individuals and pathogen to be ‘well-mixed’ throughout the network. Making this assumption allows us to derive the most common compartmental model used for environmentally-transmitted pathogens from the system of equations (13) and (14) [6, 8, 18, 37].

We use the homogeneous random variables ℐ^∗^ and 𝒫^∗^ as defined by equation (6), allowing us to simplify equations (13) and (14). Under the homogeneity assumption, we arrive at the following system

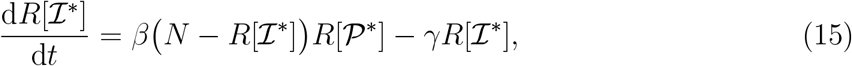

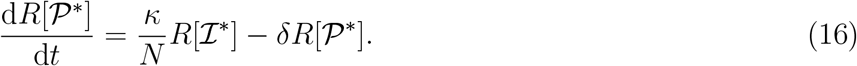

Notice that the movement parameter *µ* has completely disappeared from this system. This can be explained by carefully considering the homogeneity assumption. Under the assumption of homogeneity, the infected individuals and pathogen should be equally spread throughout the system. This means that for any two adjacent nodes, the amount of pathogen on each will always be equal, hence the net change in pathogen population caused by a movement event will be zero.

To complete our model, we must also apply this assumption to the initial conditions. The initial conditions are

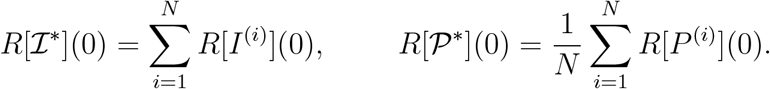

This ensures the same initial population in both the heterogeneous and homogeneous models, while removing spatial information.

We considered two different models that arise from this derivation. The first model (Model A) is a system of two equations (15) and (16). Due to the small size of the system, there are many ways in which the system can be analysed, however the homogeneity assumption means that the model does not retain any concept of the network structure. The second model (Model B) is made up of equations (13) and (14) and contains 2*N* equations (two per node). This system is more difficult to analyse but can be solved numerically with little computational expense. Since this system does not use the homogeneity assumption, it maintains some spatial aspects of the network, losing some spatial information via the independence assumption. Both models are summarised in the Appendix.

## 4 Moment closure system

In order to develop approximations with more relaxed assumptions, we begin the derivation again, this time without making the independence assumption between the infection and pathogen states of each node. As such, we retain terms of the form *ρ*_*i*_{*S*_*k*_} and *ρ*_*i*_{*I*_*k*_}. We begin by writing the differential equation for the probability of a node being infected *and* in pathogen state *k*

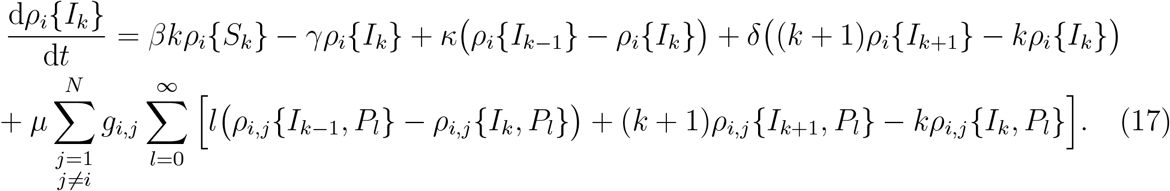

Solving the system of equations (11) and (17) will allow us to approximate the expected populations of infected individuals and pathogen, however there are still three terms with forms whose values are not immediately known, namely *ρ*_*i*_{*S*_*k*_}, *ρ*_*i,j*_{*P*_*k*_, *P*_*l*_} and *ρ*_*i,j*_{*I*_*k*_, *P*_*l*_}. The term *ρ*_*i*_{*S*_*k*_} can be calculated directly from equations (11) and (17) via the marginal probability equation *ρ*_*i*_{*S*_*k*_} + *ρ*_*i*_{*I*_*k*_} = *ρ*_*i*_{*P*_*k*_}.

The other two terms, *ρ*_*i,j*_{*P*_*k*_, *P*_*l*_} and *ρ*_*i,j*_{*I*_*k*_, *P*_*l*_}, require a choice to be made. We could choose to approximate both terms via a spatial independence assumption between nodes

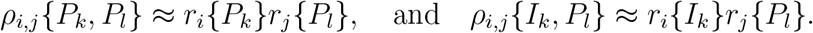

While this model would still use an independence assumption, it would only be applied between pathogen states of adjacent nodes and not between the infection state and pathogen state of each individual node. For our purposes, we would prefer to try to model the dependence of all second order terms, which should include adjacent pathogen states.

Alternatively, we could derive a differential equation for *ρ*_*i,j*_{*P*_*k*_, *P*_*l*_} (as it is a second order term) while applying a higher-order approximation to *ρ*_*i,j*_{*I*_*k*_, *P*_*l*_} (as it is a third order term). In this case, it is typical to approximate with either a pair approximation or triple correlation term, given by the following equations respectively

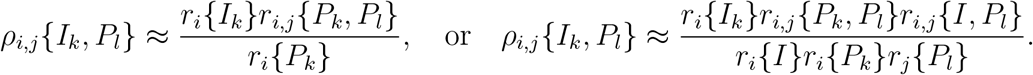

While either of these terms would result in a closed system of equations, neither allow us to write our system of differential equations in terms of expectations, leaving us with a countably infinite system of differential equations. While there are potential resolutions to the size of the system, we would prefer our solution in terms of expectations for comparison with Model A and Model B.

Instead of the above approaches, we can sum over the exact equations, equations (11) and (17), in order to write differential equations for the expected values of the exact system. Just as before, the movement terms in equation (11) simplify to first order terms when we calculate expected values. Similarly, the terms in equation (17) when summed, simplify to first and second order terms. Therefore, we can write the exact equation for the rate of change of our expected value terms without having to first construct an approximate system.

First, we take equation (17) and sum over all pathogen states, resulting in the following expressions

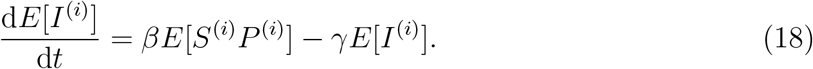

While in the previous example the term *E*[*S*^(*i*)^*P*^(*i*)^] never appeared due to the independence assumption between infection and pathogen states, here we maintain this term. As such, we require a rate of change equation for *E*[*S*^(*i*)^*P*^(*i*)^]. We note that *E*[*S*^(*i*)^*P*^(*i*)^] = *E*[*P*^(*i*)^] − *E*[*I*^(*i*)^*P*^(*i*)^], terms that we can calculate from equations (11) and (17) respectively.

Due to the nature of equation (11), we obtain the same equation for the rate of change of *E*[*P*^(*i*)^] as in the previous example

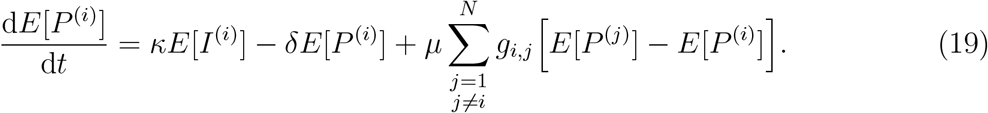

To write the differential equation for *E*[*I*^(*i*)^*P*^(*i*)^], we require the following expected values

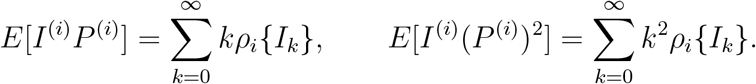

The rate of change equation for *E*[*I*^(*i*)^*P*^(*i*)^] is then obtained by multiplying equation (17) by *k* and summing over all pathogen states *k*, resulting in

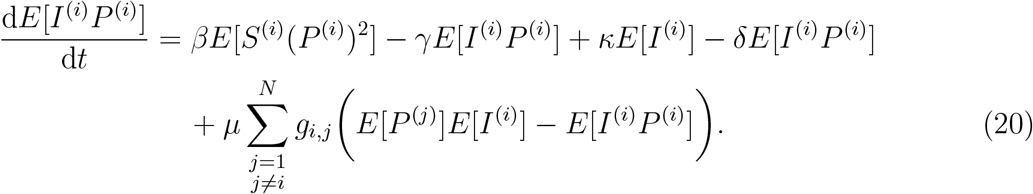

Unfortunately, in constructing the differential equation for *E*[*I*^(*i*)^*P*^(*i*)^] we have introduced a new term, namely the infection term *E*[*S*^(*i*)^*P*^(*i*)2^]. An expected value made up of the product of *n* random variables is referred to as an *n*th order moment. Hence, while all other terms in the system can be written as first and second order moments, this new term is a third order moment. If we were to write a differential equation for this term, it would require information about a fourth order moment, with *n*th order moments generally depending on the (*n* + 1)th order moment. Continuing this process indefinitely increases the size of the system. Instead, we can make a new approximation using a technique known as moment closure [22, 27, 51].

First, we look at a simple case of moment closure that was implicitly used in the previous section by comparing equation (18) to equation (13). There is a difference here in the infection term, based on whether or not the independence assumption is applied (equation (9)). This point is demonstrated by a small rearrangement of the term *E*[*S*^(*i*)^*P*^(*i*)^]. We define *µ*_*S*_ = *E*[*S*^(*i*)^] and *µ*_*P*_ = *E*[*P*^(*i*)^] and then consider the following

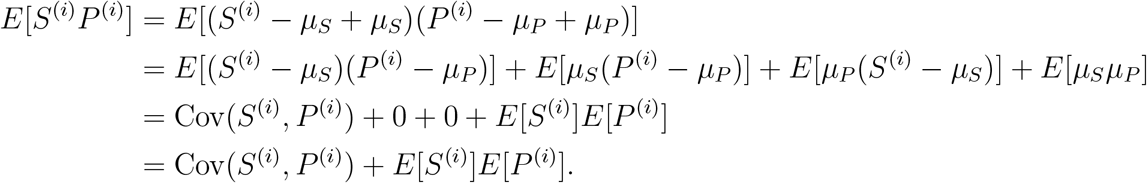

A central moment is the expectation of the product of the difference between random variables and their means. For a single random variable, the second central moment is variance, the third is skewness, etc.. For a pair of random variables, the covariance is the second central moment. To apply moment closure, we make an assumption about the distribution of the random variables, allowing us to allocate a value to the central moment.

When considering exact values, *E*[*S*^(*i*)^*P*^(*i*)^] and *E*[*S*^(*i*)^]*E*[*P*^(*i*)^] differ only by the central moment Cov(*S*^(*i*)^, *P*^(*i*)^). In the previous section, we assumed independence between the random variables *S*^(*i*)^ and *P*^(*i*)^ for all nodes *i*. When two random variables are independent, their covariance is zero, hence the independence assumption in the previous section implicitly lead to the following moment closure approximation

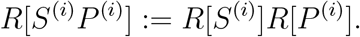

Setting Cov(*S*^(*i*)^, *P*^(*i*)^) = 0 is the simplest example of moment closure. Using moment closure allows for higher order moments to be neglected or expressed in terms of lower order moments of random variables in a system. As a result, we need only consider a finite number of moments, hence resulting in systems whose solution can be calculated.

We can use moment closure to remove all third order moments by first rewriting them in terms of their central moments. Consider the following

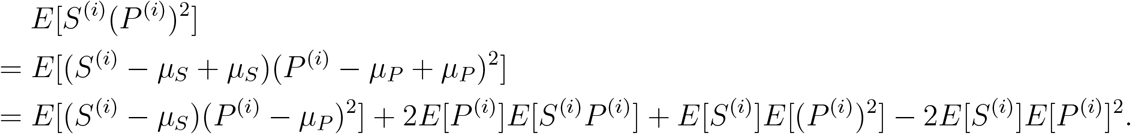

By setting the third order central moment to zero, we assume that the distributions of *S*^(*i*)^ (and hence *I*^(*i*)^) and *P*^(*i*)^ are a multivariate normal distribution [30]. We discuss the consequences of this assumption in Section 5.

We can then define an approximate system equivalent to the exact system in terms of structure and initial condition, with the following approximation applied

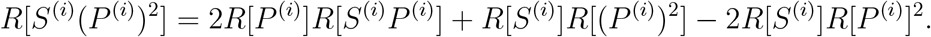

Therefore, the approximation for equation (20) is

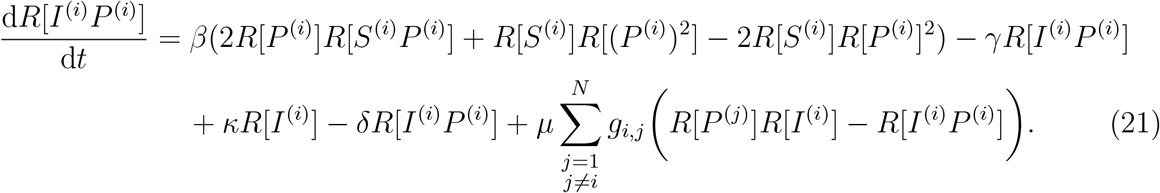

This system still requires a rate of change equation of one final term, namely *R*[(*P*^(*i*)^)^2^]. In order to obtain the rate of change equation for this term, we take equation (11) and multiply by *k*^2^ before summing over all pathogen states. The resulting equation is

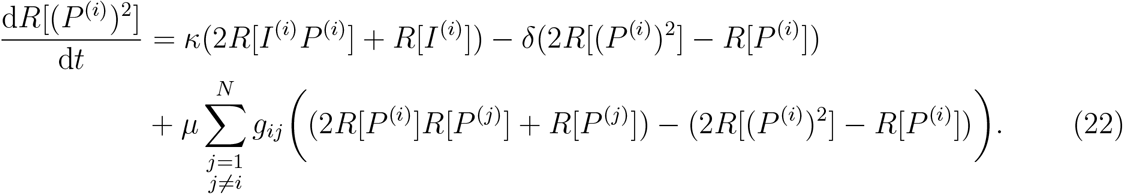

From this point we can solve the system. Any terms *R*[*S*^(*i*)^] are rewritten as 1 − *R*[*I*^(*i*)^] and then we need only numerically solve equations (21) and (22), along with the analogous approximations of equations (18) and (19) for each node *i*.

Once again, we cannot make further reductions to the approximate system without making an assumption of homogeneity. We define random variables ℐ^*∗*^ and 𝒫^*∗*^ as in the previous section, along with the random variable 𝒮^*∗*^ = *N* − ℐ^∗^.

Applying the homogeneous random variables to each equation in our approximate system yields

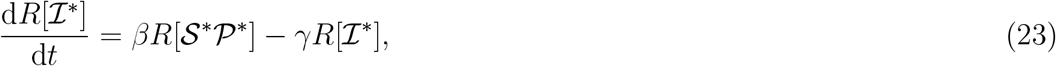

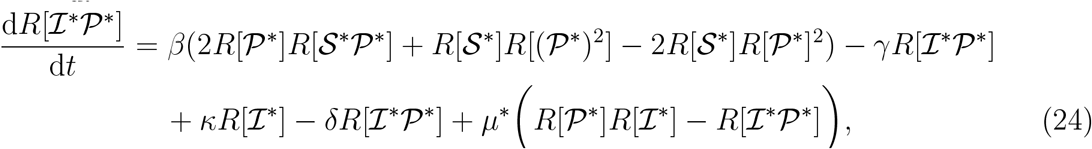

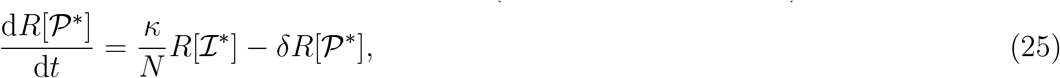

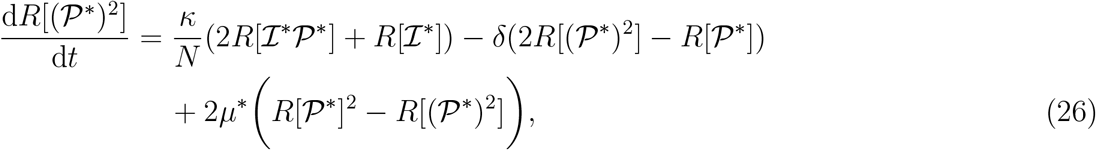

recalling that *µ*^∗^ is the adjusted movement rate given by equation (7).

Just as in the previous section, we consider two models based on this derivation. Equations (23), (24), (25) and (26) (Model C) give a system of four equations. A system of this size can be analysed, but like Model A it suffers from the restrictions of the homogeneity assumption. Model C maintains some understanding of the network structure, as is apparent by the presence of movement terms in the equations. Equations (18), (19), (21) and (22) (Model D) is a system of 4*N* equations. This is the most complicated model of those we consider, however it also comes with the fewest assumptions. We summarise Models C and D in the Appendix.

## 5 Numerical comparisons

Having derived Models A - D, we now wish to assess the performance of each model. The assessment considers both the accuracy of the model compared to the expected population of the exact system, as well as the number of differential equation in the model, a key factor in determining the computational expense of solving the system. When assessing the accuracy, expected population of the exact system cannot be calculated directly. As such, we estimate the value using the estimated average population calculated via 200 realizations.

The primary difficulty in assessing the performance of the models is the large number of changes that can be made to the system. In particular, the choice of parameters, the network structure and the choice of initial conditions can all impact the performance of the models. Below we present a summary of our findings via Figures 2 - 5 in which we explore the models while considering the above changes. We remind the reader that the goal of this paper is the derivation of the models, hence our exploration of the models is primarily qualitative.

**Figure 2:**
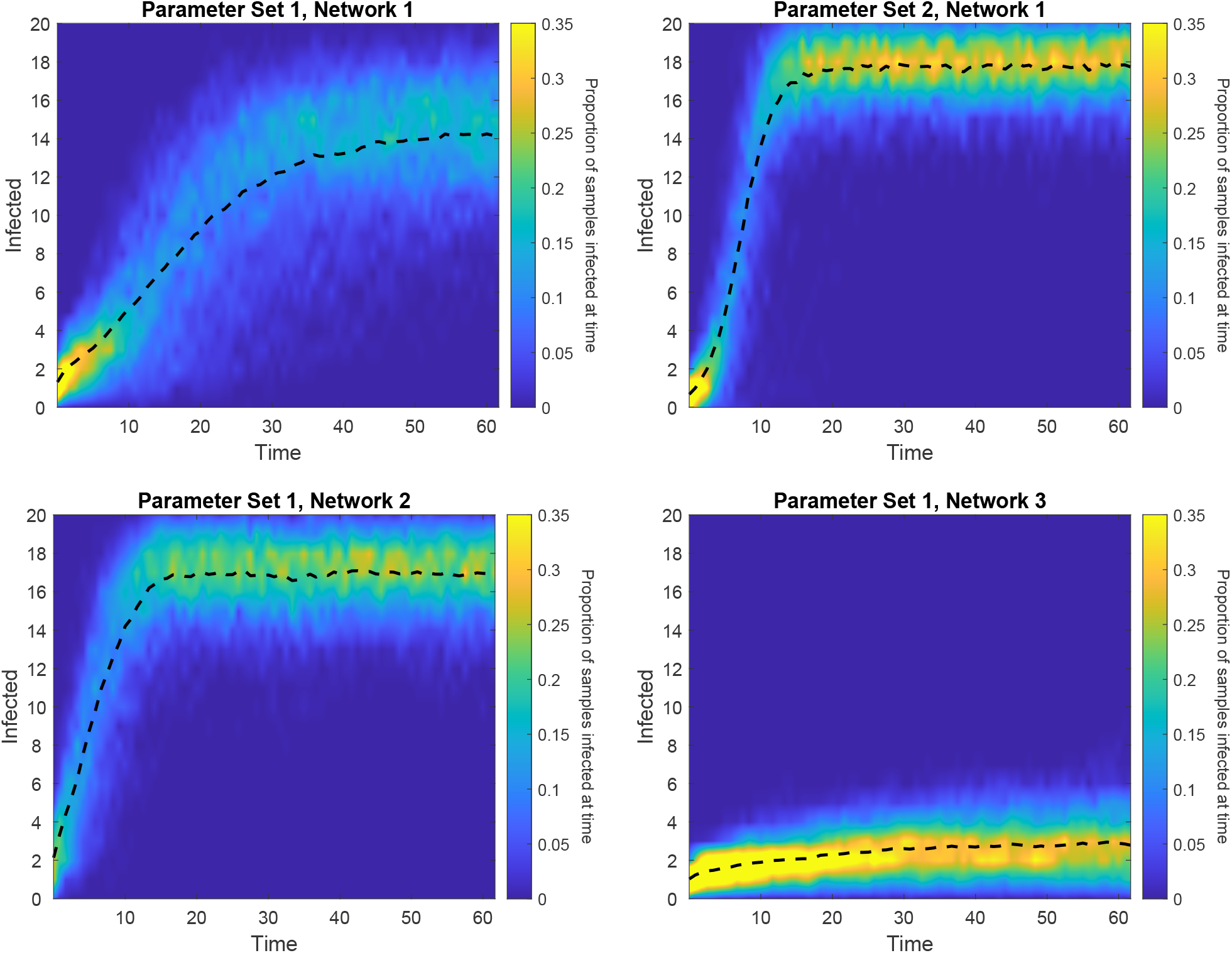
Heat-maps showing the spread of the data collected from the 200 realizations. Specifically, the colour at a given coordinate represents the proportion of the 200 realizations that have a given number of infected individuals at that time. The black dashed line shows the estimated average population over the 200 realizations. Each figure uses the movement rate *µ* = 0.1. The corresponding network and parameter set (see Section 5.2) are stated above each figure.

### 5.1 Model recap

Before comparing our models, we quickly recap the assumptions applied to each model as summarised in Table 2. Models A and B are derived under an independence assumption between the pathogen state and the infection state of each node. Models C and D bypass this independence assumption, but as a result require the use of a moment closure assumption, assuming that for each node, the pair of random variables *I*^(*i*)^ and *P*^(*i*)^ are multivariate normally distributed. Models A and C are derived under the homogeneity assumption, meaning that all nodes are indistinguishable, while Models B and D are heterogeneous, fully taking into account the structure of the network and individual properties of each node.

**Table 2:**
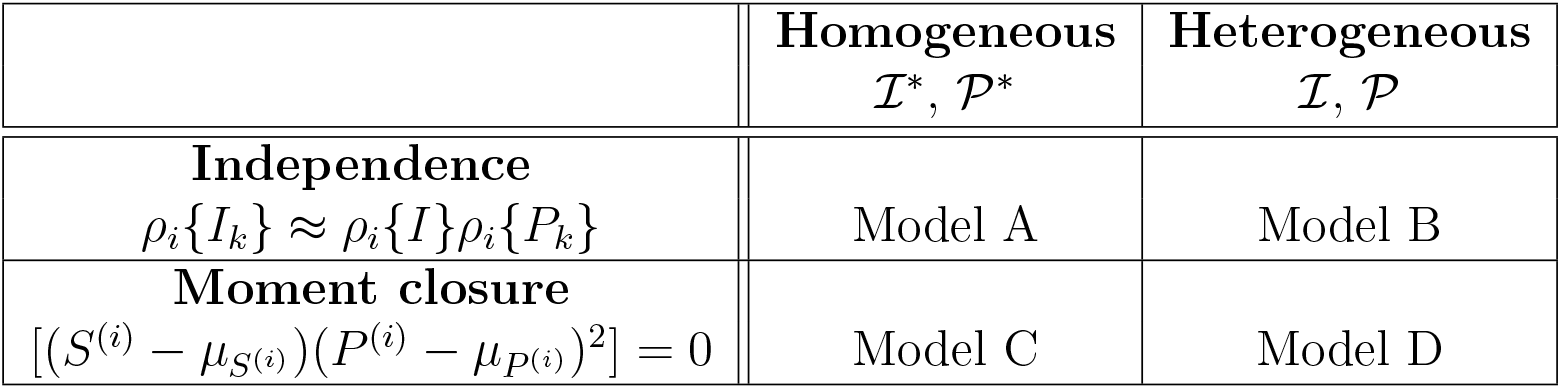
Summary of assumptions applied to each model.

| | **Homogeneous**<br>$\mathcal{I}^*, \mathcal{P}^*$ | **Heterogeneous**<br>$\mathcal{I}, \mathcal{P}$ |
| --- | --- | --- |
| **Independence**<br>$\rho_i\{I_k\} \approx \rho_i\{I\}\rho_i\{P_k\}$ | Model A | Model B |
| **Moment closure**<br>$[(S^{(i)} - \mu_{S^{(i)}})(P^{(i)} - \mu_{P^{(i)}})^2] = 0$ | Model C | Model D |

Model A enforces the most restrictive set of assumptions, while Model D enforces the most relaxed set of assumptions. Therefore, we expect to observe Models B - D outperforming Model A in every scenario. With Model D having the most relaxed assumptions, we expect it to be the closest approximation to the exact system and hence outperform Models A - C. The purpose of this analysis is to test the models’ performance with respect to each other, to observe the magnitude of improvement that comes from less restrictive assumptions and to test the accuracy of the models relative the estimated averages population.

### 5.2 Parameter choices and network structures

In order to compare the approximations of our models to the simulated network, we require choices of both parameter sets and network structures. We will explore the models over two parameter sets (see Table 3) and over a variety of networks, all established below.

**Table 3:**
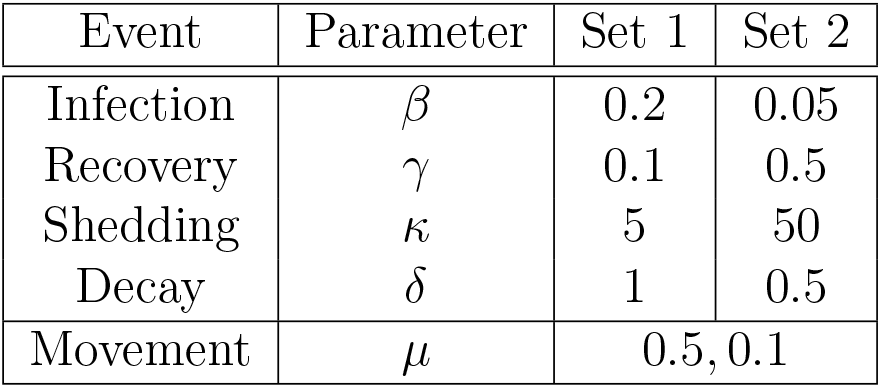
Parameter values used for the different rates in the network. We consider two different parameter choices: Set 1 and Set 2. For each parameter set, we consider two different movement rates.

Parameter set 1 was chosen to have similar scales of shedding and decay rates, *κ* and *δ*, resulting in a low level of pathogen being present in the network at any given time, hence less potential for it to spread. As fewer pathogen units are present, we have a higher infection rate *β* and lower recovery rate *γ* (relative to parameter set 2) to ensure that the infection does not die out. Parameter set 2 has a much larger shedding rate *κ* than decay rate *δ*, resulting in the potential for pathogen to survive for relatively long times, hence allowing it more time to travel between nodes. As there is significantly more pathogen in the system, we considered a lowered infection rate *β* and a heightened recovery rate *γ* (relative to parameter set 1) while still maintaining a reasonable population. For each parameter set, we also explore two different movement rates *µ* to observe the impact of spreading the pathogen through the system at different speeds.

For our network structure, we will be considering four different networks, each with *N* = 20 nodes. Network 1 is randomly constructed such that each node has an average of five edges per node. This is achieved by randomly allocating 5*N/*2 edges between nodes. There is also a secondary condition that the network must be connected (there exists a set of edges connecting each pair of nodes). The other networks are constructed to consider extreme cases. Network 2 is fully-connected, with edges between each pair of nodes (*g*_*ij*_ = 1 for all *i* ≠ *j*). Network 3 is a chain, with nodes in a line and edges between successive nodes (*g*_*ij*_ = 1 iff | *i* − *j*| = 1). Network 4 has all nodes exclusively adjacent to a central node (*g*_*ij*_ = 1 iff *i* = 1, *j* ≠ 1).

Finally, we note that the above parameters and networks are those being presented, but not the only choices that we considered and analysed. The choices presented allow us to demonstrate interesting and meaningful behaviour that we observed throughout our analysis. Throughout our research we considered numerous different networks including increasing the number of nodes, random networks with varying densities, and other extreme cases like Networks 2 - 4. We also observed numerous parameter sets, in particular focusing on how the movement rate changed the results.

### 5.3 Simulated system details

Before getting into our comparison, we address some of the details that are necessary to ensure the success of the simulated system. Realizations of the CTMC are implemented using the algorithm described in Section 2.2.2. We average our results over 200 realizations, in order to ensure that the estimated average population accurately represents the expected population of the exact system. Figure 2 shows a heat-map of the results of the 200 realization at each time, demonstrating the distribution of the samples.

The other important detail of the simulated system is the choice of initial condition. We have run our model with no initially infected individuals, one node containing 30 pathogen units and no pathogen on any other nodes. That is

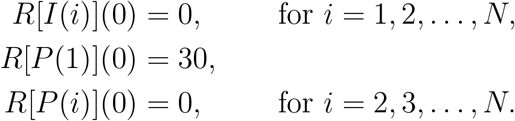

The primary importance of the initial condition is to avoid extinction events which can cause discrepancies when comparing systems based on whether they treat the population as discrete or continuous. When the population is zero in a simulated realization, there is the potential for the pandemic to end if the pathogen is unable to infect an individual before it all decays. However, due to the continuous nature of expectations used in the model, the expectation of a fraction of an individual can prevent extinction from occurring. As such, we wish to avoid initial conditions that would cause the population to die out early (for instance, starting with a single infected individual).

### 5.4 Results and discussion

#### 5.4.1 Parameter choices

We begin by observing the effects of changing the parameters with a fixed network structure (namely Network 1). We will first consider Figure 3, in which we compare the models and simulated realizations where the movement rate is higher, specifically *µ* = 0.5, and then consider Figure 4, in which the comparison is made with a lower movement rate, specifically *µ* = 0.1.

**Figure 3:**
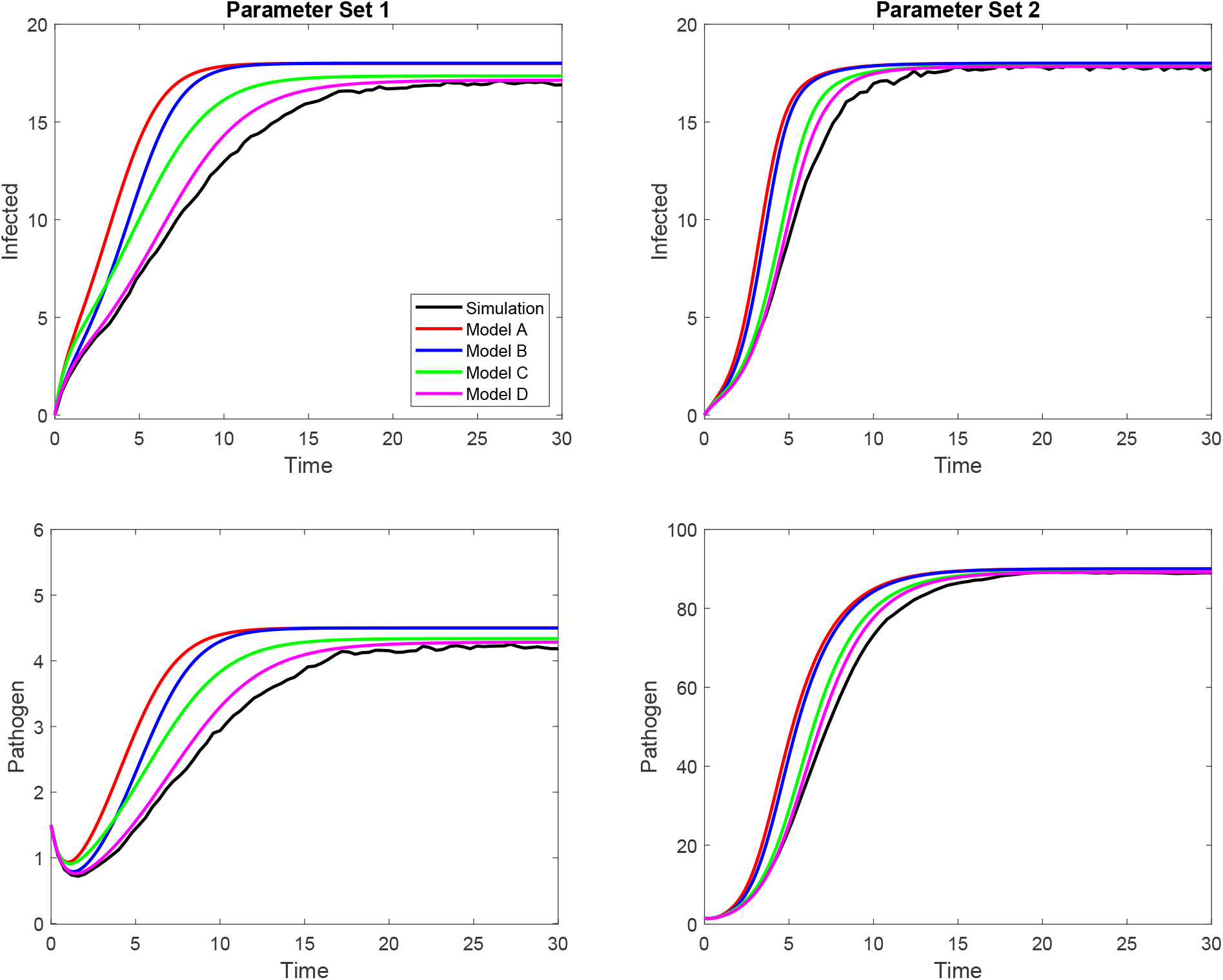
Comparison between Models A - D (colored lines) and the estimated average population over 200 realizations (black line) for ‘higher’ movement rate *µ* = 0.5 over a random network with 20 nodes (Network 1 in Section 5.2). Parameter Sets 1 and 2 are shown on the left and right respectively (see Table 3) with infected populations shown in upper plots and pathogen populations shown in the lower plots. We observe that the more restrictive assumptions produce a greater error between the estimated average population and each model, with Model D giving a relatively accurate approximation.

**Figure 4:**
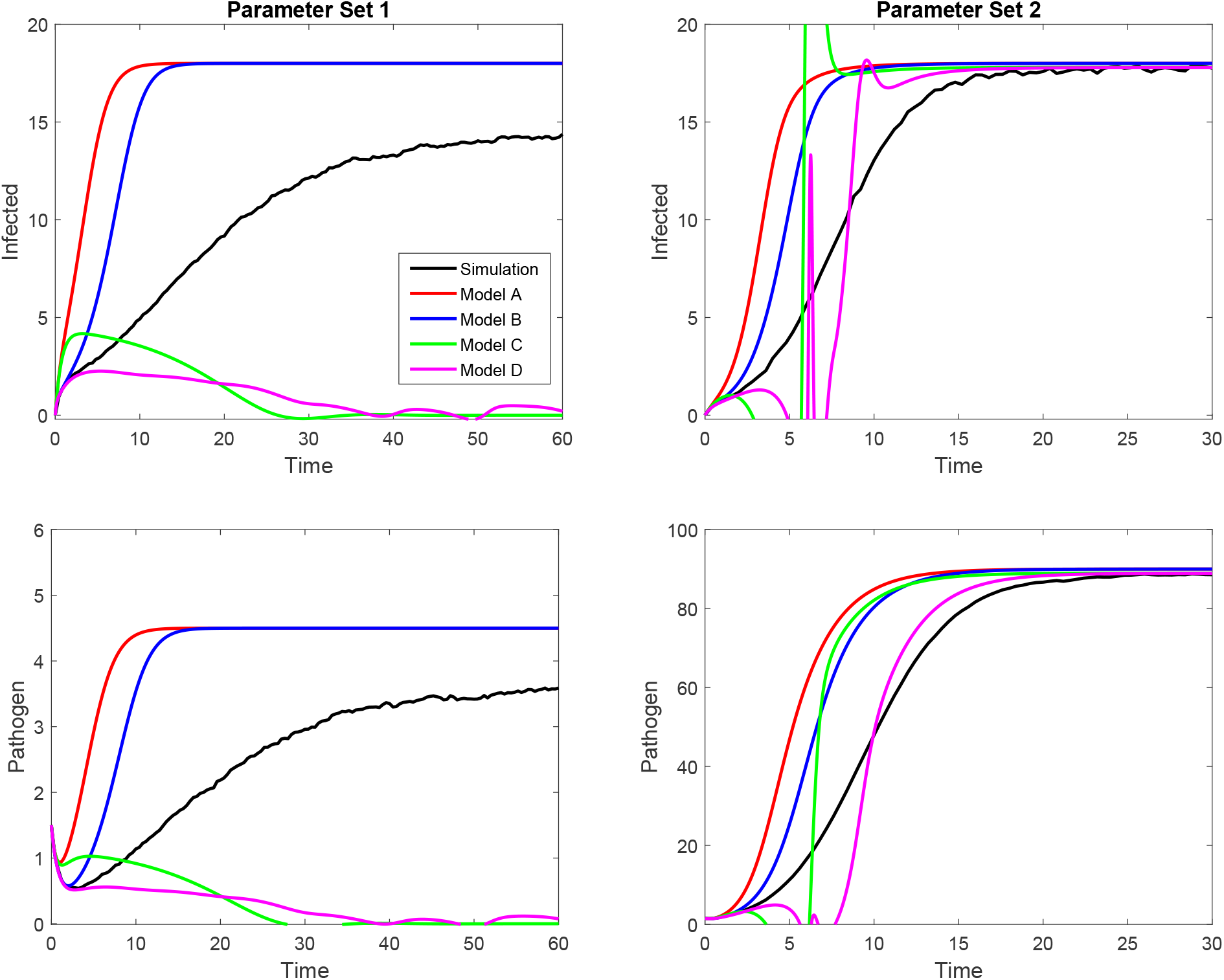
Comparison between Models A - D (colored lines) and the estimated average population over 200 realizations (black line) for ‘lower’ movement rate *µ* = 0.1 over a random network with 20 nodes (Network 1 in Section 5.2). Parameter Sets 1 and 2 are shown on the left and right respectively (see Table 3) with infected populations shown in upper plots and pathogen populations shown in the lower plots. We observe divergent, non-physical behaviour from Models C and D, while Models A and B are stable but give extremely inaccurate approximations.

In Figure 3 we observe exactly what we had anticipated. Model D is the closest approximation to the estimated average population, vastly outperforming Model A. For every choice of parameters we considered, we observed this same outcome of Model D outperforming Model A, so long as the movement rate *µ* is ‘sufficiently high’ (this movement rate caveat will be discussed below when considering Figure 4).

We also observe Models B and C performing somewhere in between Models A and D. Note that for parameter set 1, Model B briefly outperforms Model C before moving back into the expected hierarchy. This observation is not too surprising as it is not immediately clear that our independence assumption compared to our moment closure assumption is a more drastic restriction than the homogeneous assumption compared to the heterogeneous assumption (see Table 2).

The other observation between the two parameter sets in Figure 3 is that there is a visible difference in the long term populations predicted by the models in parameter set 1, but not in parameter set 2. While not shown here, we observed that the distinction between the long term predicted populations for parameter set 1 disappeared when the movement rate *µ* became sufficiently large. In general, we saw that for any choice of parameters, if the movement rate is increased sufficiently all models will yield the same long term result. This tells us that the error in both the homogeneity assumption and the independence/moment closure assumptions will become zero as *µ* → ∞. This result is expected as sufficiently high movement rates cause homogeneous mixing among the population.

In Figure 4 we observe that both Models C and D are having stability issues, yielding physically unreasonable results. By running the models over numerous movement rates, we observed that the instability occurs for *µ <* 0.13 for parameter set 1 and *µ <* 0.44 for parameter set 2. For all parameter sets that we tested, if the movement rate becomes ‘sufficiently low’ then we observe this unstable behaviour in Models C and D. Given that this issue only occurs for Models C and D, we conclude that the issue arises due to the moment closure approximation. We consider this issue further when considering different network structures below.

In circumstances where Models C and D show divergent behaviour we observed the transient behaviour of Models A and B to be extremely inaccurate approximations to the estimated average population. In fact, we will show in Section 5.4.3 that the unstable behaviour of Models C and D will only occur when Models A and B are extremely inaccurate approximations.

From our observations, Model D appears to be a relatively accurate approximation for the estimated average population in cases where it remains stable. In cases where the movement rate is extremely high, all the models give the same predictions with Model A being the most efficient (smallest number of differential equations). In cases where the movement rate is sufficiently low, Models C and D show divergent behaviour and Models A and B are extremely inaccurate approximations.

#### 5.4.2 Network structures

Next we consider observing the effects of changing the network structure with a fixed parameter set (namely parameter set 1 with *µ* = 0.1). This comparison is shown in Figure 5 using the four different networks established in Section 5.2. The impact on the accuracy and stability of the models across different networks is relatively independent of the choice of parameter set (beyond what has already been discussed in Section 5.4.1) hence only one parameter set is considered here.

**Figure 5:**
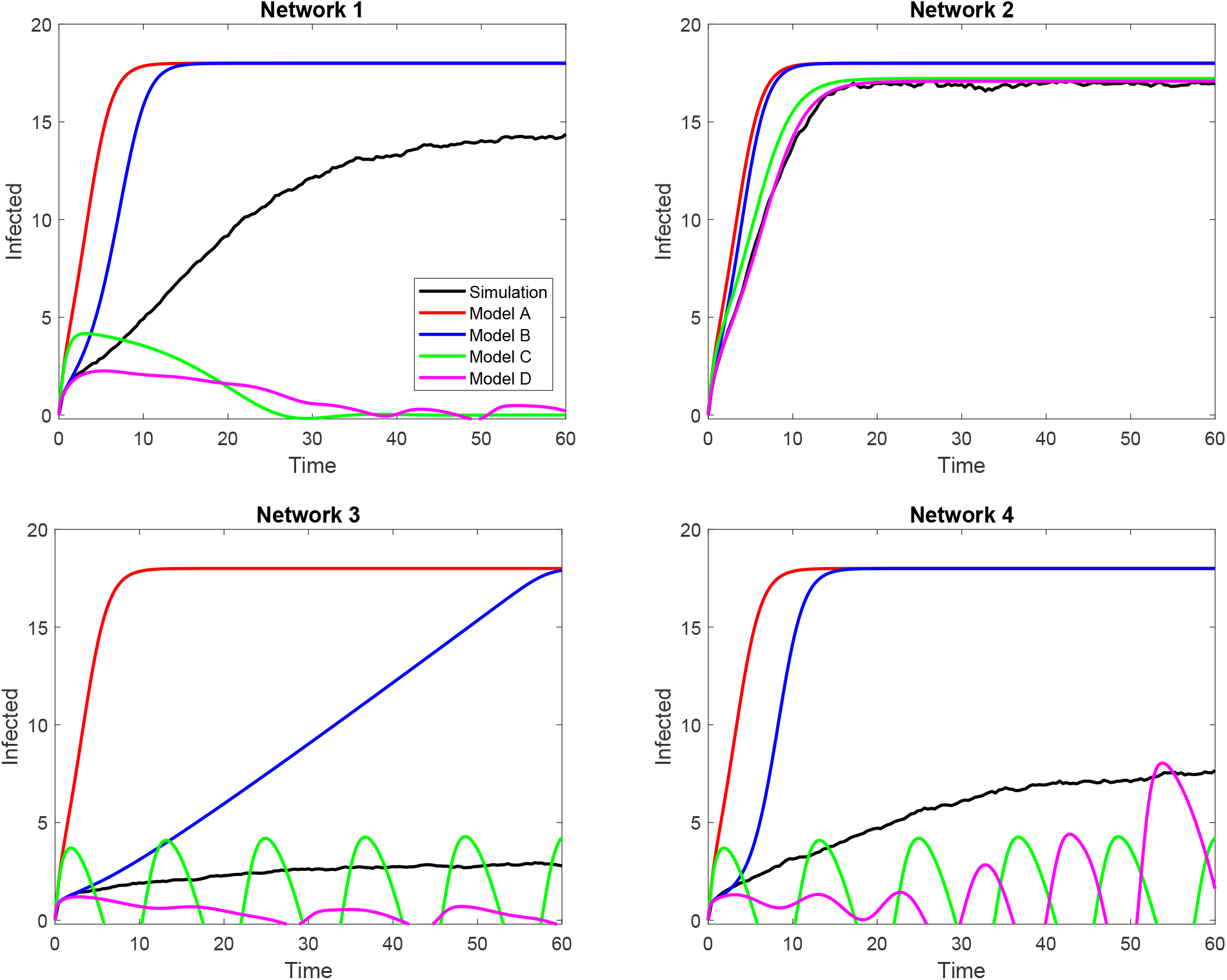
Comparison between infected populations for Models A - D (colored lines) and the estimated average population over 200 realizations (black line) for parameter set 1 with *µ* = 0.1 (see Table 3) over four different network structures (see Section 5.2). Networks with more edges between nodes increase the movement rate of pathogen through the network, and hence produce more accurate and stable results.

Network 1 in Figure 5 was already considered in Section 5.4.1 but is useful here as a baseline for our extreme network structures. The fully-connected Network 2 yields the best approximations, with Models C and D demonstrating stable behaviour and Model D being an extremely accurate approximation to the estimated average population. Further, every model gives its most accurate approximation for this network. There are two clear reasons for improved approximations when considering Network 2. The first is that the network is closer to satisfying the homogeneous assumption, as all nodes are connected to each other. The second is that the movement rate (despite being fixed at *µ* = 0.1) is artificially increased. The movement rate is defined per edge, hence for a pathogen unit on a node with *M* edges, the true movement rate is *Mµ*. We have already seen that an increase movement rate results in stable behaviour of these models.

While Network 2 is somewhat ideal for approximating, we also want to consider extreme cases that are the worst case for approximating. Both Networks 3 and 4 have approximately the same number of edges (one per node) but different structures. In these cases, the small number of edges means that the true movement rate is much less than in the random network case. As such, the unstable behaviour of Models C and D in these cases is unsurprising.

We also note the drastic difference given by Model B in Networks 3 and 4, which can be understood by the difference in their network structures. In Network 3, for a pathogen unit to move from node 1 to node 20 would require it to move along the chain through all other nodes. In Network 4, pathogen at node 1 can immediately move to any other node. As a consequence of these structures, pathogen takes longer to spread through Network 3, hence slowing the infection rate in this network, while pathogen can move to all nodes immediately in Network 4, hence speeding up the infection rate. This difference is only recognized by the heterogeneous models, Models B and D, as they recognize the distinction between populations on nodes (although this is not visible for Model D due to the instability). Similarly, Model C cannot identify the structure due to the homogeneity assumption, but gives different results based on the number of network edges. As such, Model C gives almost identical results when calculating an approximation for the populations of Networks 3 and 4, despite their structural differences. Finally, Model A does not take into account any network structure, including the number of edges. As such, it gives identical results for all four networks.

#### 5.4.3 Issues with the closure approximation

The most important conclusion to be drawn from the above figures is that Model D yields accurate approximations for the expected population of the exact system in cases where the model is stable. We aim now to gain a further understanding of the unstable nature of Models C and D. To do this, we considering the moment closure assumption and the distributions of *I*^(*i*)^ and *P*^(*i*)^.

Our moment closure assumption is that *I*^(*i*)^ and *P*^(*i*)^ are multivariate normally distributed random variables (for each node *i*). There are a number of reasons that this assumption is extremely inaccurate. Considering the possible values of the random variable *I*^(*i*)^ and that individuals can only be infected or susceptible, there is no way that any sort of normal distribution would be a suitable approximation.

In contrast to *I*^(*i*)^, the random variable *P*^(*i*)^ can take on a variety of integer values, giving it potential to be approximated by a binomial distribution and hence a normal distribution. However, this too has an issue when we consider the absorbing zero state of the system, that is when the network contains no infected individuals and no pathogen. Given our initial condition, it is not infrequent that the infection simply dies out before secondary infections occur. As such, the distribution of *P*^(*i*)^ is generally bimodal, with a set of values that form a peak that could be approximated by a normal distribution and a second peak given by all the zero values resulting from the absorbing state.

These issues prevent the moment closure approximation from being sufficiently accurate, and although there are a number of potential resolutions, they each come with their own difficulties. If we restructure the model, preventing the zero state from occurring (by say setting all rates into the zero state to zero), then we would resolve the primary issue with the distribution of *P*^(*i*)^ [21, 38, 39]. Unfortunately, we would still have our issue with the random variable *I*^(*i*)^. Given that *I*^(*i*)^ is a random variable with two outputs (susceptible or infected), it is most likely represented by some hypergeometric distribution. We could attempt a closure approximation by combining a multivariate hypergeometric and normal pair of random variables (for *I*^(*i*)^ and *P*^(*i*)^ respectively), however finding the moment-generating function for this distribution would be extremely difficult or even impossible, as it is not guaranteed that the moment-generating function exists.

There is another potential resolution in which we apply the moment closure approximation later in the derivation upon reaching the population random variables ℐ and 𝒫. If we prevent the zero state issue, then both of these distributions independently are likely to appear as approximately normally distributed. However, at this point, we have already applied the homogeneity assumption, which as discussed earlier in the context of Model C, is likely to prevent accurate approximations. Further, while the distributions of each random variable may appear as normal when observed separately, we expect relatively strong correlation between the number of infected individuals and the pathogen count in the network. Therefore, even if we resolved all of these issues, there’s no guarantee that the approximation would become substantially more accurate.

To finish our discussion, we discuss the behaviours for low movement rates in the context of the moment closure approximation. Consider the following expectation that is approximated by the moment closure assumption

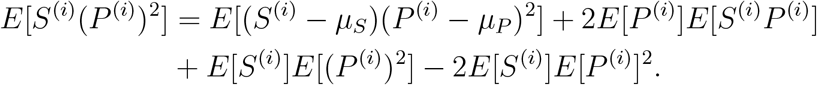

In particular, we take an interest in the following terms

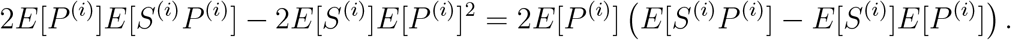

The term in brackets here is precisely the difference in the infection term between the true solution and Model B. Based on the difference between the average simulated estimate and Model B in Figure 4, we observe that there is a consistent result that

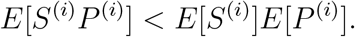

Most importantly, this difference becomes greater as the movement rate *µ* decreases, observable by the increasing distance between the average simulated estimate and Model B. Therefore, we expect that the following is generally true

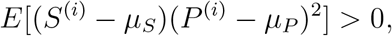

and that this prevents the infection term, specifically *E*[*S*^(*i*)^(*P*^(*i*)^)^2^], from becoming negative. When we set the third order central moment to zero via the moment closure approximation, the new term *R*[*S*^(*i*)^*P*^(*i*)2^] now has the potential to become negative, particularly when *µ* is small. In cases where *R*[*S*^(*i*)^*P*^(*i*)2^] becomes negative, we obtain a variety of physically unreasonable results (as seen in Figures 4 and 5). To resolve this issue, we could instead consider a different form of closure approximation, for instance

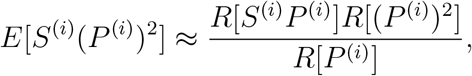

or something proportional to this [23, 26]. An approximation term is always positive, preventing the negativity issue above. Although it may look somewhat different to the approximation terms used in our current approximation, an assumption of independence applied to both approximations will reduce to the same expression. The exact derivation of such a term is somewhat more complicated, hence we leave this for future work.

## 6 Conclusion

Development of more detailed ODE models and understanding of the underlying assumptions are necessary steps if we wish to draw meaningful conclusions when modeling the spread of infections. This is particularly important when considering environmentally-transmitted diseases, due to the significance of spatial structures inherently present in the physical system. Our approaches demonstrate how techniques used in the direct-transmission literature can be applied to environmentally-transmitted pathogens, specifically assumptions of independence and higher order moment closure approximations. Using these techniques has allowed us to rigorously present the derivation of a system of ODEs whose structure mirrors the compartmental ODEs frequently used to model environmental transmission in mathematical epidemiology, and to present other potential candidate models that may yield improved results.

A primary difficulty moving from directly-transmitted infections to environmentally-transmitted pathogens is the representation of the pathogen population. By clearly describing the dynamics of the individual-based network model, we have demonstrated that approximate systems of differential equations can be rigorously constructed, yielding the corresponding rate of infection term and the pathogen growth term. This also fundamentally relied on a clear definition of pathogen units and their interactions with susceptible and infected individuals.

Detailed derivations of the approximate models and their comparison to simulated realizations of the network model allowed us to qualitatively measure the consequences of different assumptions. Comparisons over a variety of network structures and parameter sets demonstrated that, in the absence of unstable behaviour, models that relaxed the assumptions of independence and spatial homogeneity yielded more accurate approximations over the average behaviour of the individual-based network model. In these stable cases, the presence of a basic independence assumption (Models A and B) caused large errors in the approximations, while the homogeneity assumption (Models A and C) caused errors that were substantial for some parameter sets. Model D, in which the heterogeneity of the system was modeled and the moment closure approximation was applied to third order terms, was able to outperform the other models in cases where the solution was stable, but at the expense of a larger system of differential equations. These results align with our expectations and the use of these types of approximations in the existing direct-transmission literature.

The improved approximations given by Model D are limited to cases where the behaviour of the solution remains stable. Slow movement rates, which in our case were due to both the choice of movement rate parameter *µ* and the sparsity of the network, caused the infection rate to become negative, resulting in physically unreasonable solutions when using Models C and D. We showed that this was a consequence of the choice of a multivariate moment closure and also argued that this would only occur when Models A and B were also yielding inaccurate approximations. It should be noted that more appropriate moment closure approximations can both prevent this unstable behaviour and further improve the accuracy given by Model D. For instance,

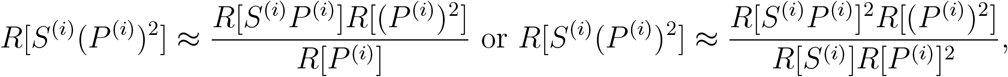

although both are more difficult to motivate than the multivariate moment closure approximation presented. Our use of the multivariate normal distribution as the moment closure approximation better aligns with our goals of presenting a clear derivation of our ODE models and the potential pitfalls of applying different approximations.

## Acknowledgements

This work was supported by the joint DMS/NIGMS Mathematical Biology Program through National Institutes of Health (NIH) award R01GM113239 and the Centers for Disease Control and Prevention (CDC) award U01CK000587.

## Appendix: Summary of ODE models: Models A - D

**Model A** s given by equations (15) and (16), and the initial conditions given below

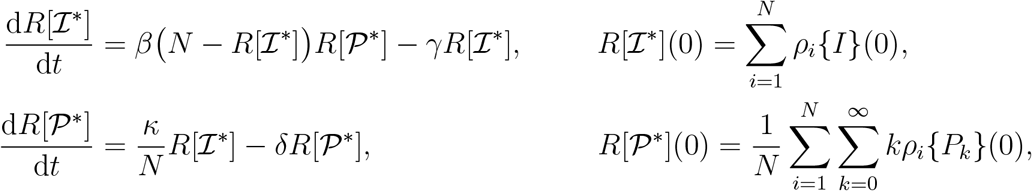

where *R*[ℐ^∗^] and *R*[𝒫^∗^] are approximations for *E*[*I*] and *E*[*P*] respectively.

**Model B** is given by equations (13) and (14), and the initial conditions given below

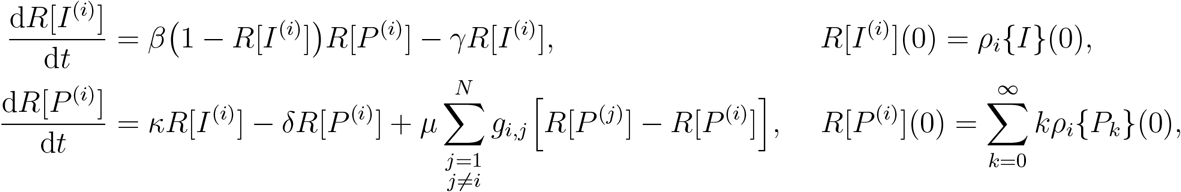

for all nodes *i* = 1, 2, …, *N*. We calculate *R*[ℐ] = ∑_*i*_ *R*[*I*^(*i*)^] and *R*[𝒫] = ∑_*i*_ *R*[*P*^(*i*)^]*/N* as approximations for *E*[ℐ] and *E*[𝒫] respectively.

**Model C** is given by equations (23), (24), (25), (26), and the initial conditions given below

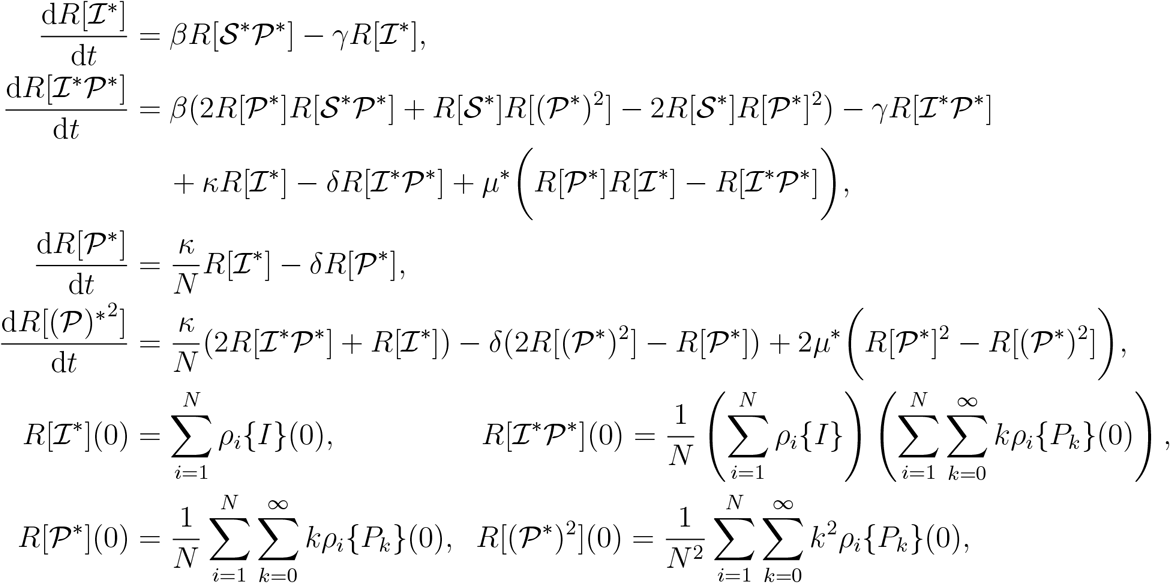

where *R*[ℐ^∗^] and *R*[𝒫^∗^] are approximations for *E*[ℐ] and *E*[𝒫] respectively.

**Model D** is given by equations (18), (19), (21), (22), and the initial conditions given below

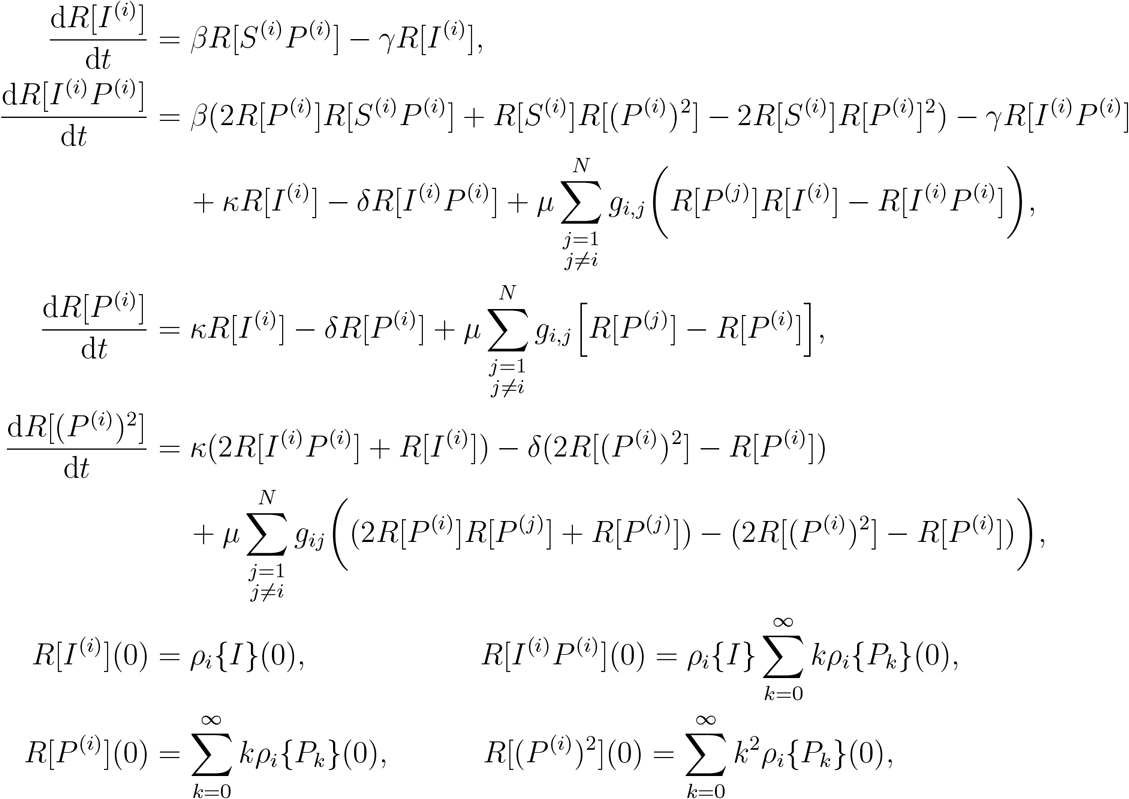

for all nodes *i* = 1, 2, …, *N*. We calculate *R*[ℐ] = ∑_*i*_ *R*[*I*^(*i*)^] and *R*[𝒫] = ∑_*i*_ *R*[*P*^(*i*)^]*/N* as approximations for *E*[ℐ] and *E*[𝒫] respectively.

